# Metabolic mechanisms of nitrogen substrate utilisation in three rhizosphere bacterial strains investigated using quantitative proteomics

**DOI:** 10.1101/627992

**Authors:** Richard P. Jacoby, Antonella Succurro, Stanislav Kopriva

## Abstract

Nitrogen metabolism in the rhizosphere microbiome plays an important role in mediating plant nutrition, particularly under low inputs of mineral fertilisers. However, there is relatively little mechanistic information about which genes and metabolic pathways are induced by rhizosphere bacterial strains to utilise diverse nitrogen substrates. Here we investigate nitrogen substrate utilisation in three taxonomically diverse bacterial strains previously isolated from Arabidopsis roots. The three strains represent taxa that are consistently detected as core members of the plant microbiome: Pseudomonas, Streptomyces and Rhizobium. We use phenotype microarrays to determine the nitrogen substrate preferences of these strains, and compare the experimental results versus computational simulations of genome-scale metabolic network models obtained with EnsembleFBA. Results show that all three strains exhibit generalistic nitrogen substrate preferences, with substrate utilisation being well predicted by EnsembleFBA. Using label-free quantitative proteomics, we document hundreds of proteins in each strain that exhibit differential abundance values following cultivation on five different nitrogen sources: ammonium, glutamate, lysine, serine and urea. Proteomic data show that the three strains use different metabolic strategies to utilise specific nitrogen sources. One diverging trait appears to be their degree of proteomic flexibility, with *Pseudomonas* sp. *Root9* utilising lysine nutrition via widespread protein-level alterations to its flexible metabolic network, whereas *Rhizobium* sp. *Root491* shows relatively stable proteome composition across diverse nitrogen sources. Our results give new protein-level information about the specific transporters and enzymes induced by diverse rhizosphere bacterial strains to utilise organic nitrogen substrates.

**Importance:** Nitrogen is the primary macronutrient required for plant growth. In contemporary agriculture, the vast majority of nitrogen is delivered via mineral fertilisers, which have undesirable environmental consequences such as waterway eutrophication and greenhouse gas production. There is increasing research interest in designing agricultural systems that mimic natural ecosystems, where nitrogen compounds are cycled between plants and soil, with the mineralisation of recalcitrant soil organic-N molecules mediated via microbial metabolism. However, to date there is little mechanistic information about which genes and metabolic pathways are induced by rhizosphere bacterial strains to metabolise organic-N molecules. Here, we use quantitative proteomics to provide new information about the molecular mechanisms utilised by taxonomically diverse rhizosphere bacterial strains to utilise different nitrogen substrates. Furthermore, we generate computational models of bacterial metabolism from a minimal set of experimental information, providing a workflow that can be easily reused to predict nitrogen substrate utilisation in other strains.

## Introduction

Improved nitrogen management in agricultural systems is crucial for environmental sustainability. Large-scale application of mineral nitrogen fertilisers has extensive off-target effects, such as greenhouse gas production and waterway eutrophication (1). One potential pathway to boost agricultural sustainability involves substituting mineral fertilisers with organic nutrients derived from recycling various waste streams. For low-input agricultural systems to provide sufficient bioavailable nitrogen to meet the demands of plant growth, future crop management practices will need to better incorporate microbial pathways of nitrogen mobilisation (2). One specific suggestion involves engineering the rhizosphere microbiome to promote the mineralisation of organic nitrogen, coupled with engineering of plant root metabolism to release rhizodeposits that recruit beneficial microbial strains (3). However, the ability to manipulate plant-microbe cooperation is limited by an incomplete knowledge of the specific microbial traits involved in root colonisation and nutrient mobilisation (4).

Nitrogen flows in the rhizosphere are complex, with plants and microbes potentially cooperating but sometimes competing for uptake of diverse nitrogen molecules (5). Legume-Rhizobia symbioses provide an example of cooperation, whereby the majority of the plant’s nitrogen nutrition is derived from bacterial fixation of atmospheric N_2_ (6). Outside of legumes, it is generally accepted that plants obtain the majority of their nitrogen nutrition from inorganic forms such as NO_3_ and NH_4_, whereas microbes are more adept at acquiring more recalcitrant organic nitrogen forms such as proteins and amino acids (7). Therefore, cooperative nutrient transfers can occur when microbes take up soil-bound organic nitrogen, which is subsequently transferred to plants in a mineralised form following microbial lysis or protozoic predation (8). Conversely, competitive flows can occur when microbes immobilise inorganic nitrogen, or when plants take up organic nitrogen (9). Adding further complexity, plant root exudates contain large amounts of organic nitrogen molecules which can serve as carbon and nitrogen substrates for bacterial growth. The rate of amino acid release from plant roots increases under exposure to specific bacterial metabolites (10), but organic nitrogen molecules released via root exudation can also be efficiently re-acquired by the root system (11).

Investigations of how bacteria utilise diverse nitrogen substrates have been documented since the beginning of modern microbiology (12). Ammonium is the preferred nitrogen source for most bacteria, and experimental designs usually include ammonium as a control treatment, to compare against alternative nitrogen sources or starvation treatments (13). Over decades, such studies have provided detailed insight into fundamental physiological mechanisms such as the molecular pathways of bacterial nitrogen assimilation, the perception of nitrogen status, and the response to nitrogen starvation in *E. coli* (14). However, other bacterial taxa possess different mechanisms for regulating nitrogen metabolism (15, 16), with soil bacteria exhibiting extensive diversity regarding their nitrogen substrate preferences and also the metabolic pathways used to metabolise organic nitrogen sources (17, 18). Therefore, novel insights into metabolic mechanisms of nitrogen metabolism may be observed by studying nitrogen substrate utilisation in taxonomically diverse bacterial strains isolated from the rhizosphere. The rhizosphere microbiome has attracted increasing research attention over the past 20 years. From the results of 16S pyrosequencing studies, it has become increasingly apparent that the rhizosphere hosts a taxonomically diverse bacterial microbiota, which plays an important role in determining plant growth and health (19). Recently, multiple research groups have established large collections of bacterial strains isolated from field-grown plants, which can be used to dissect the functional traits carried out by individual strains, or reassembled into synthetic communities that recapitulate microbiome function (20, 21). There is now an opportunity to study these plant-associated microbial strains using high-throughput ‘omics techniques, to acquire new insights into the specific molecular mechanisms that confer a selective advantage in the plant-associated niche (22).

Alongside experimental approaches, computational modelling is becoming a widespread approach to investigate microbial metabolism (23). One particularly useful method is the construction of genome-scale metabolic network models, which translate the information encoded in the bacterial genome into a computational formalism that can be analysed with mathematical methods (24). However, curated genome-scale metabolic models are only available for a relatively small set of extensively studied bacterial strains, and generally it is difficult to analyse newly sequenced bacterial strains using computational modelling. This limitation exists because reconstructing a curated genome-scale metabolic network model is a painstaking process that requires extensive manual curation as well as the acquisition of devoted experimental data, particularly regarding biomass composition. Although progress is being made towards automated reconstruction of genome-scale metabolic network models, many challenges still have to be addressed (25). Recently, a method named EnsembleFBA has been proposed as a potential approach to approximate genome-scale metabolic networks for diverse bacterial strains. Instead of relying on the availability of a single manually curated genome-scale model, EnsembleFBA uses the information derived from multiple metabolic networks, which are reconstructed from the same initial draft network and refined through the process of positive and negative gapfilling on randomized sets of growth and non-growth conditions (26). As a proof of concept, it was shown that the EnsembleFBA method achieved greater precision in predicting essential genes than an individual, highly curated model.

Here we investigate nitrogen metabolism in three taxonomically diverse bacterial strains previously isolated from Arabidopsis roots. We apply a combination of methods, including quantitative proteomics, growth assays, phenotype microarray and EnsembleFBA. With the proteomic data, we were particularly interested in determining the specific proteins that are enriched according to different nitrogen sources, to decipher the metabolic strategies used for nitrogen acquisition across different rhizosphere bacterial strains. In parallel, we applied the EnsembleFBA method to reconstruct and analyse sets of genome-scale metabolic network models for each strain, using the phenotype microarray data for training and testing the model predictions of nitrogen substrate utilisation.

## Results

We studied nitrogen metabolism in three taxonomically diverse bacterial strains isolated from roots of field-grown Arabidopsis: *Pseudomonas* sp. *Root9*, *Streptomyces* sp. *Root66D1* and *Rhizobium* sp. *Root491*. Strains were previously isolated in Bai et al (20), and the three strains chosen here correspond to taxa that were repeatedly observed to be highly abundant in the microbiome of field-grown Arabidopsis plants (20, 27, 28).

### Measurement and modelling of growth phenotypes on different nitrogen sources

First, we investigated each strain’s ability to utilise 94 diverse nitrogen sources using a phenotype microarray (BIOLOG PM3B) (Supplementary Figure 1, Supplementary Table S1). The data reveal that all three strains can catabolise a relatively high number of substrates, with the three strains exhibiting positive growth phenotypes on 55-61 of the 94 substrates tested. This indicates that all three strains have generalistic nitrogen substrate preferences, which has been previously suggested to be a selective advantage in the rhizosphere (29). In parallel, we used EnsembleFBA (26) to test how accurately nitrogen substrate utilisation can be computationally predicted across the three strains (Supplementary Figure 1, Supplementary Table S1). When nitrogen substrate utilisation is assessed in binary terms (growth versus no growth), there is a good concordance between the experimental results and the computational predictions, with Ensemble FBA showing an accuracy in predicting growth in about 78% of cases for the three strains (Supplementary Table S2). However, there is a relatively poor correlation between the proxy values of metabolic activity predicted by the models versus the experimental measurements, with a comparison of percentile rank between the datasets yielding r^2^ values between 0.23 and 0.5 across the three strains (Supplementary Figure S2). Interestingly, the accuracy of the model prediction seems to vary across different molecular classes, with good concordance for amino acids but poor concordance for nitrogen bases (Figure 1).

**Figure 1:**
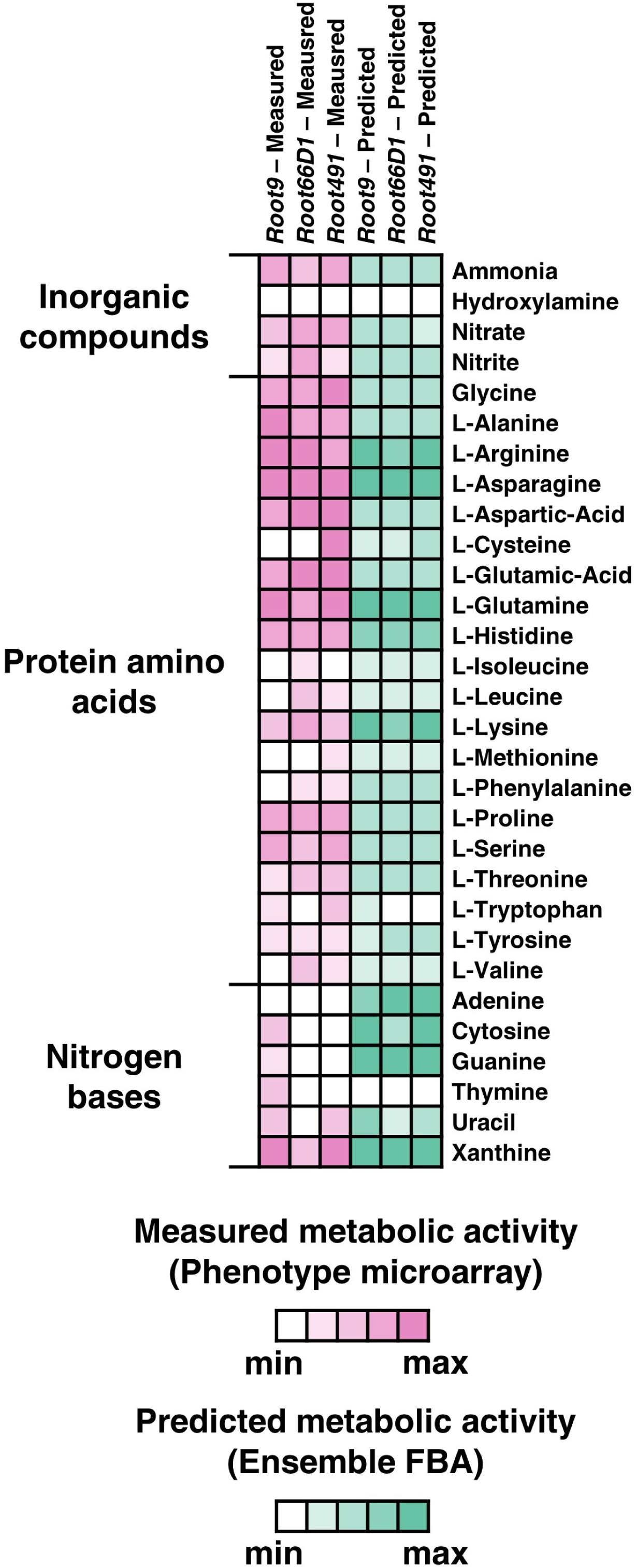
Nitrogen substrate preferences of three rhizosphere bacterial strains assessed via Phenotype Microarray and EnsembleFBA. Displayed here are results for 30 nitrogen substrates selected from the 94 tested. White boxes indicate no metabolic activity, whereas boxes with darker shades correspond to higher metabolic activity, either measured via Phenotype Microarray (pink) or predicted via EnsembleFBA (green). Metabolic activity values were z-score normalised within each strain.

### Growth curves in batch culture

We conducted growth curves in batch culture to further investigate the growth phenotypes of these three strains when cultivated on five selected nitrogen sources (ammonium, glutamate, lysine, serine and urea), (Figure 2, Supplementary Table S3). The rationale for selecting these nitrogen sources is because ammonium serves as the inorganic reference, the three chosen amino acids are abundant in soils and exhibit diverse charges (glutamate negative, lysine positive, serine neutral), and urea is a widely applied agricultural fertiliser. Nitrogen concentration in the medium was 5 mM, which was determined to be a yield-limiting nitrogen concentration in all three strains (30), (Supplementary Figure S3). In Pseudomonas sp. *Root9*, we see that lysine nutrition elicits a long extension of the lag phase (Figure 2A), perhaps indicative of a physiological reprogramming that must occur before rapid proliferation can proceed (31). In contrast, *Rhizobium* sp. *Root491* exhibited very similar growth curves across all five nitrogen sources, indicative of growth homeostasis across different nutrient sources.

**Figure 2:**
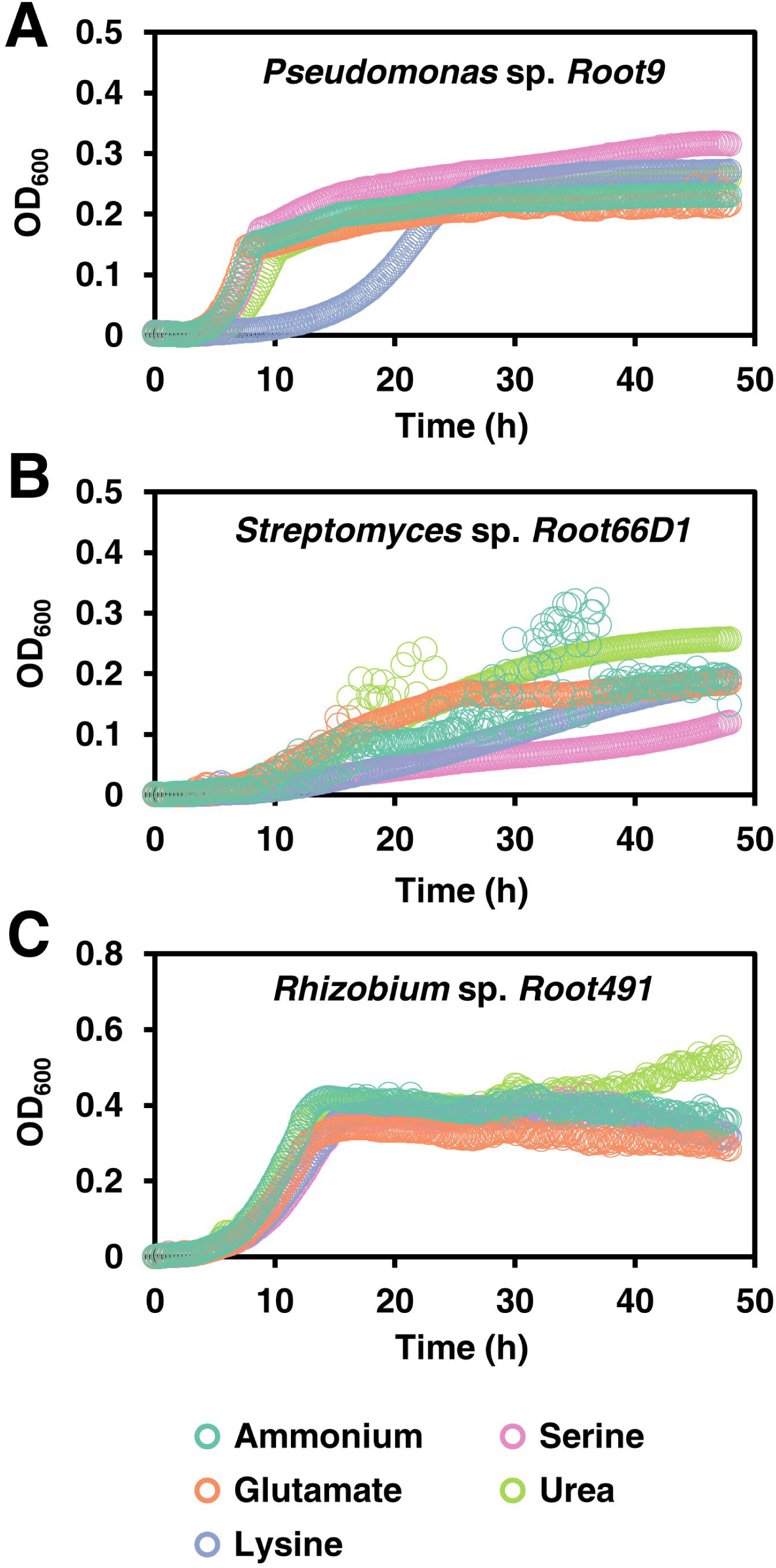
Growth curves of three rhizosphere strains cultivated on five nitrogen sources. Cultures were grown in 48-well plates on minimal medium containing a single nitrogen source. OD_600_ (uncorrected for path length) was logged every 10 min using a plate reader.

### Proteome remodelling in response to different nitrogen sources

The main aim of this study was to define systems-level differences in cellular proteome composition in three rhizosphere bacterial strains cultivated on five different nitrogen sources. Therefore, bacteria were cultivated on the same nitrogen sources shown in Figure 2 (ammonium, glutamate, lysine, serine and urea), cells were harvested during the exponential growth phase, and cellular protein composition analysed using label-free quantitative proteomics. A numerical summary of protein IDs is shown in Table 1, a visual overview of the derived results is shown in Figure 3, volcano plots for all 10 pairwise comparisons across all three strains are shown in Supplementary Figures S4-S6, and the MaxQuant abundance values for all detected proteins are given in Supplementary Table S4.

**Figure 3:**
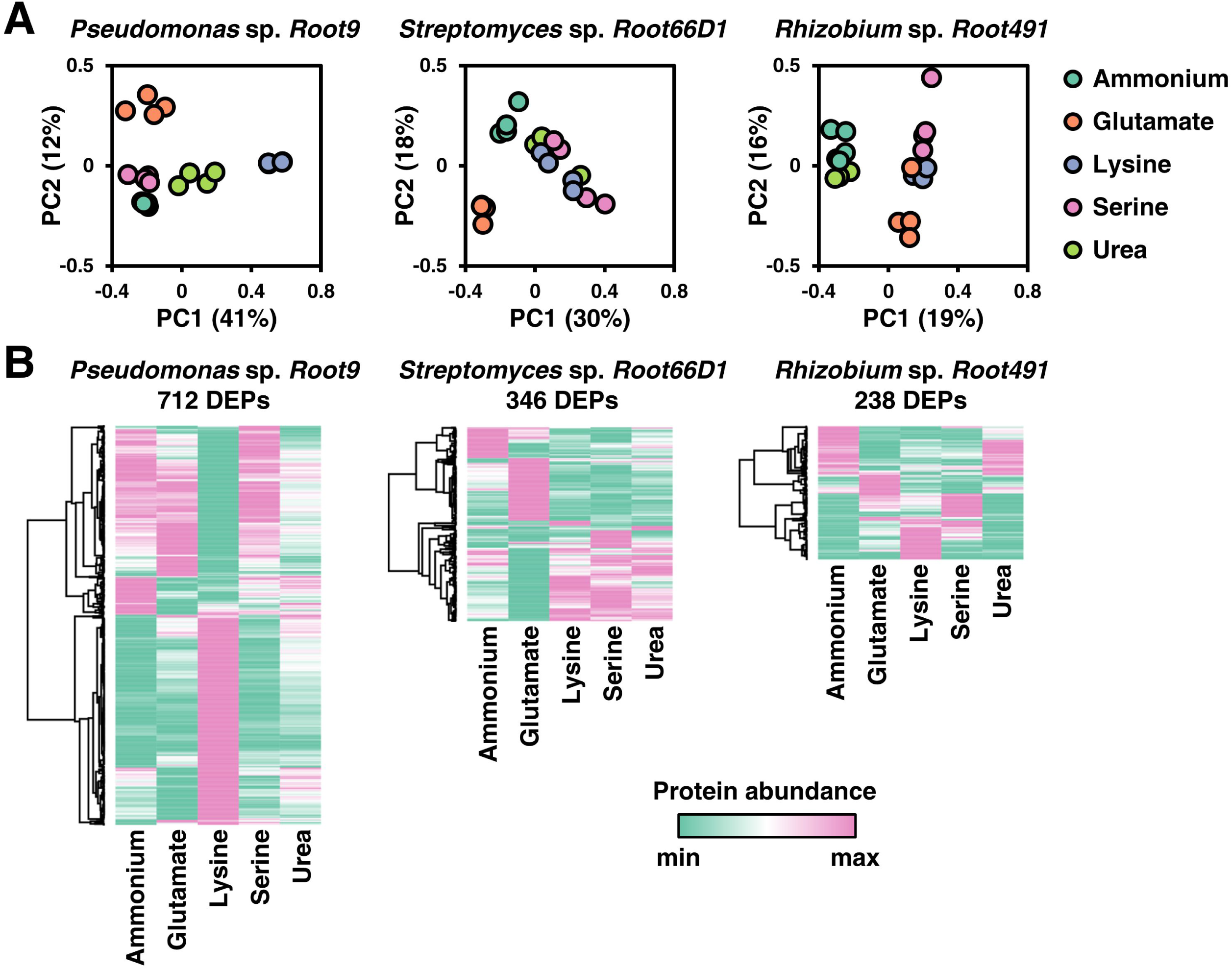
Overview of proteome composition in three rhizosphere bacterial strains when cultivated on five nitrogen sources. A: Principal component analysis (PCA) of the five different nitrogen sources for each strain. B: Heat maps of protein abundance for differentially expressed proteins (DEPs) for each of the three strains. To define DEPs, protein abundance in one condition was compared to its abundance in the other four conditions. If in any of these 10 comparisons, a protein has a log2FC > 1 and a BH-p-value < 0.05, then it is considered a DEP. Only DEPs that were detected in at least three replicates for all five nitrogen treatments are included in the heatmaps. Rows were clustered using Pearson’s correlation coefficient.

**Table 1:**
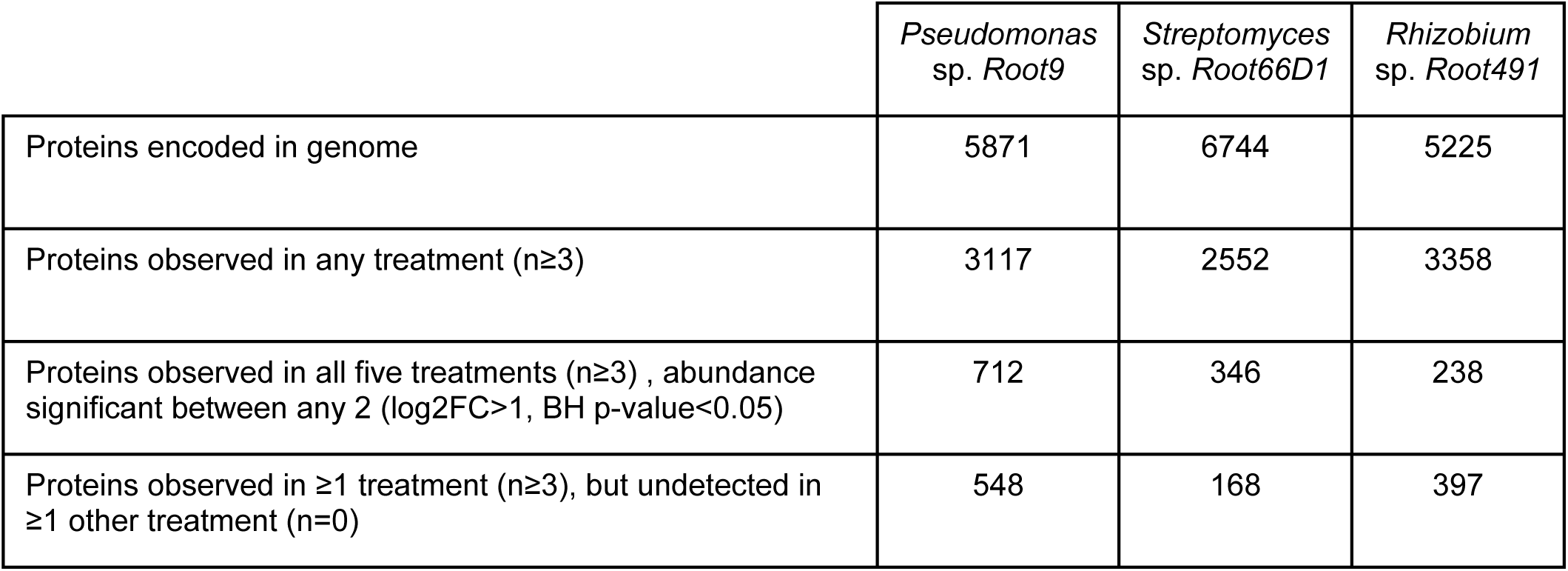
Summary of label free quantitative proteomic data for three rhizosphere bacterial strains cultivated on five different nitrogen sources.

|  | *Pseudomonas*<br>sp. Root9 | *Streptomyces*<br>sp. Root66D1 | *Rhizobium*<br>sp. Root491 |
| --- | --- | --- | --- |
| Proteins encoded in genome | 5871 | 6744 | 5225 |
| Proteins observed in any treatment (n≥3) | 3117 | 2552 | 3358 |
| Proteins observed in all five treatments (n≥3) , abundance significant between any 2 (log2FC>1, BH p-value<0.05) | 712 | 346 | 238 |
| Proteins observed in ≥1 treatment (n≥3), but undetected in ≥1 other treatment (n=0) | 548 | 168 | 397 |

Comparing protein composition across the three strains, it seems that *Pseudomonas* sp. *Root9* exhibits more protein-level flexibility compared to the other two strains. This is evident in the PCAs and heatmaps presented in Figure 3, which show that lysine treatment of *Pseudomonas* sp. *Root9* elicits a large proteomic remodelling compared to the other four nitrogen treatments, characterised by hundreds of differentially expressed proteins. In contrast, we see that *Rhizobium* sp. *Root491* exhibits a degree of proteomic homeostasis across the different nitrogen treatments, as shown by the closer clustering of the PCA data points and the lower number of differentially expressed proteins in this strain.

Comparing across the five different nitrogen sources, we see that each individual nitrogen source seems to elicit a differential proteomic impact in the three different strains. For example, lysine nutrition elicits large-scale changes in the proteome of *Pseudomonas* sp. *Root9*, yet relatively few proteomic changes in the other two studied strains. In both *Streptomyces* sp. *Root66D1* and *Rhizobium* sp. *Root491*, urea nutrition elicited no proteomic changes compared to ammonium, whereas in *Pseudomonas* sp. *Root9* there were over 100 proteins with differential abundance values between ammonium versus urea (Supplementary Figures S4-S6).

### Orthologous proteins and metabolic pathways modulated by nitrogen nutrition

To allow inter-strain comparisons of the label-free quantitative proteomic data acquired from the three taxonomically diverse rhizosphere bacterial strains, we utilised cross-species gene annotation via KEGG orthologues (32). We selected individual proteins that represent the 495 KEGG orthologues which were detected in all five treatments across all three strains, and visualise the abundance of these representative orthologues using a heatmap and PCAs in Figure 4, with numerical data provided in Supplementary Table S5. As can be seen in Figure 4A and 4B, the samples group together according to the three bacterial strains rather than the five nitrogen sources. This indicates that the baseline differences in strain-specific proteome composition are much greater than any treatment-induced differences elicited by nitrogen nutrition. In Figure 4C we plot a PCA of these 495 KEGG orthologues when protein abundance in the four organic nitrogen sources is normalised versus the inorganic nitrogen source ammonium. This shows that lysine nutrition in *Pseudomonas* sp. *Root9* elicits a proteomic response that is qualitatively different compared to the strain-medium combinations profiled in this study.

**Figure 4:**
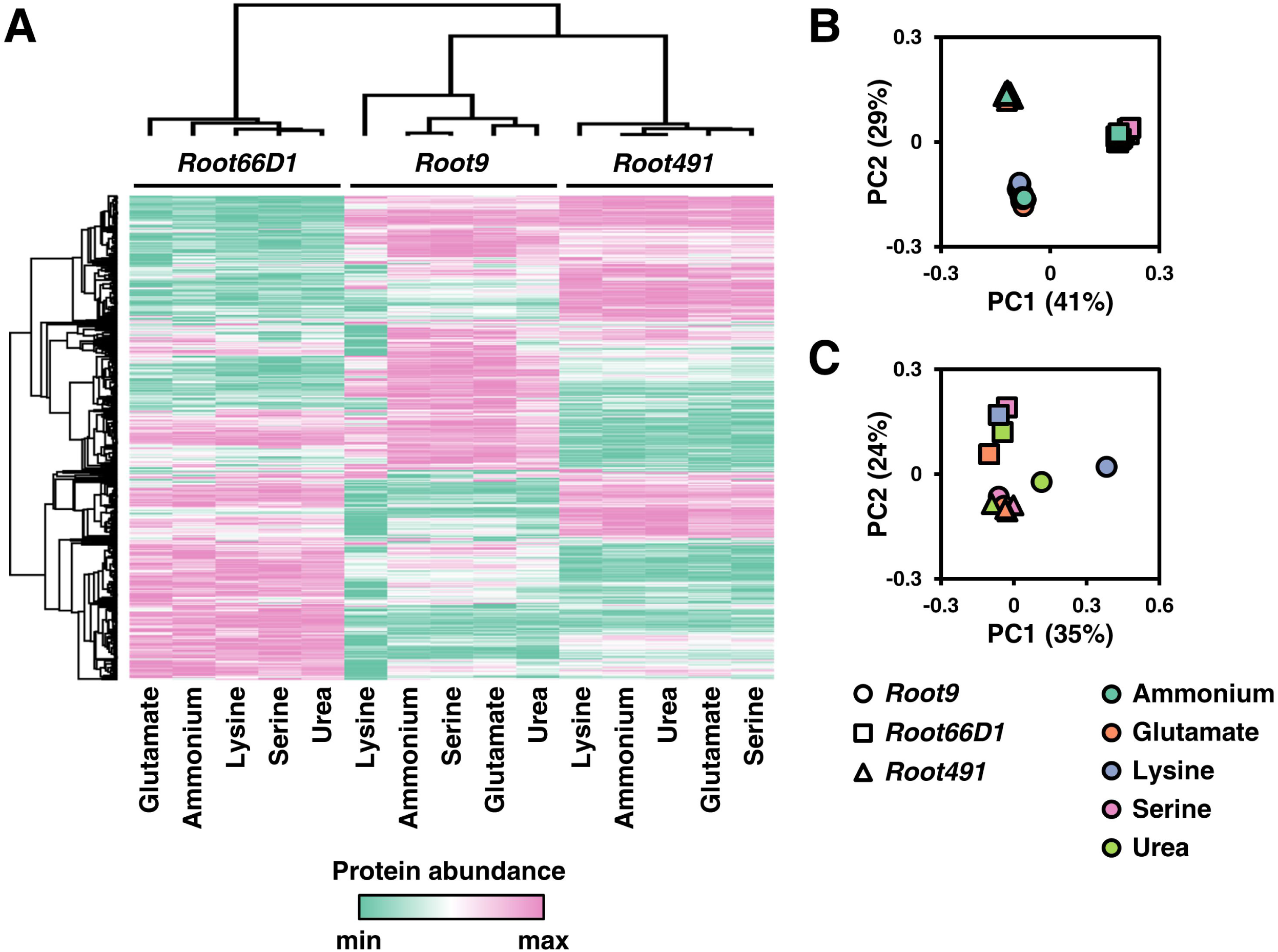
Comparison of protein abundance values for 495 KOs (Kegg orthologs) across three rhizosphere bacterial strains cultivated on five nitrogen sources. A: Heat map of KO abundance across the three rhizosphere bacterial strains cultivated under five nitrogen sources. B: Principal component analysis (PCA) of KO abundance across the three rhizosphere bacterial strains cultivated under five nitrogen sources. C: Principal component analysis (PCA) of KO abundance across the three rhizosphere bacterial strains for the four organic nitrogen sources, when KO abundance was normalised to ammonium (inorganic reference). The KOs annotated to proteins via IMG were matched across the proteomic dataset for the three bacterial strains. Data was filtered to contain only the 495 KOs that were observed in all four replicates across all five treatments in all three strains. MaxQuant LFQ abundance values were z-score normalised within each strain. Rows and columns were clustered using Pearson’s correlation coefficient.

Our next step was to analyse which specific KEGG pathways were modulated according to nitrogen treatment in the three strains. In Figure 5, we show the results of Fisher’s exact test to determine whether the constituent proteins of 30 KEGG pathways exhibited altered abundance profiles in the 10 pairwise comparisons between different nitrogen sources. Numerical data for all 126 tested pathways compared is provided in Supplementary Table S6. Looking at the specific pathways modulated by nitrogen nutrition across the three strains, it seems that *Rhizobium* sp. *Root491* undergoes fewer alterations to KEGG pathways related to metabolism, but instead exhibits extensive modulation to pathways related to environmental processing and motility. For *Pseudomonas* sp. *Root9* and *Streptomyces* sp. *Root66D1*, we see that many of the pairwise comparisons are characterised by widespread modulation to all KEGG pathways, indicating that extensive proteome remodelling has taken place between the different nitrogen sources.

**Figure 5:**
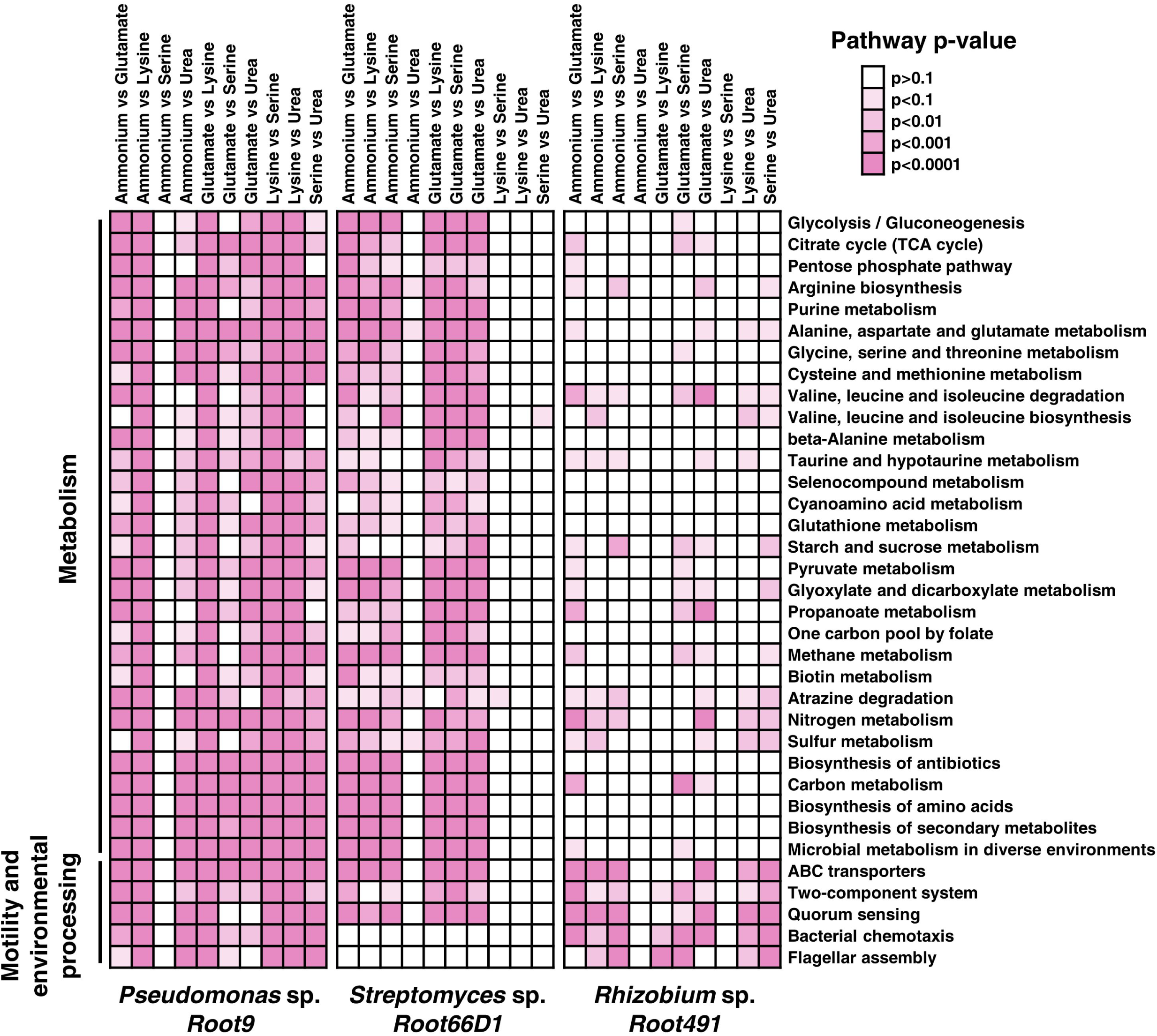
Assessment of KEGG pathways that were modulated at the protein abundance level between different nitrogen treatments. Kegg orthologs annotated to proteins via IMG were matched to KEGG pathways, and Fisher’s exact test was used to determine the statistical significance of pathway modulation between two nitrogen treatments. Darker shades of pink represent lower p-values via Fisher’s exact test. Pathways with fewer than three identified proteins were excluded from analysis. This figures shows the 30 pathways with the highest number of significantly differences between treatments (p<0.01), data for all ∼100 pathways are in Supplementary Table S6.

Next, we compared the metabolic flux distributions outputted from EnsembleFBA versus the differentially expressed proteins identified in the quantitative proteomic datasets (Supplementary Tables S7-S13). To visualise how nitrogen source affects protein abundance and computationally predicted fluxes, we used the Interactive Pathway Explorer to map KEGG orthologues and reactions onto the KEGG map ‘Metabolic Pathways’ (33). Visualisations for each of the three amino acid treatments (glutamate, lysine and serine) in pairwise comparisons versus ammonium were produced for both the proteomic data (Supplemental Figure S7) and also the computational modelling data (Supplemental Figure S8). Overall, it is evident that a similar set of metabolic pathways have been mapped in both the experimental and computational approaches, with good coverage of glycolysis, TCA cycle, and amino acid metabolism. However, there is relatively little concordance between the differentially regulated metabolic steps identified by the proteomics data versus the differentially regulated fluxes outputted by EnsembleFBA. For instance, the proteomic data show that lysine nutrition elicits significant modifications to lipid metabolism in *Pseudomonas* sp. *Root9*, whereas many of the reaction steps in lipid metabolism are absent from the EnsembleFBA flux distributions. This difference could derive from a known limitations of genome-scale modelling approaches such as EnsembleFBA, because we used a generic biomass function to construct the models, which does not account for variations in bacterial lipid composition between genotypes and treatments (34). Therefore, improved model accuracy probably requires condition-specific measurement of microbial biomass composition.

### Proteins correlated to the PII protein of the nitrogen stress response

Analysing the quantitative proteomics data, we noticed that the different nitrogen sources often elicited changes in the abundance of proteins involved in the well-characterised nitrogen stress response, such as GlnK (PII protein), amtB (ammonium transporter) and GlnA (glutamine synthetase) (14). Therefore, we postulated that our dataset may allow us to discover new proteins that are regulatory targets of the nitrogen stress response in less studied bacterial taxa. We first analysed the abundance of PII, a well characterised protein of the nitrogen stress response that exhibited significantly different abundance values between certain nitrogen treatments in all three strains (Figure 6A). Next, we assessed which other proteins in the dataset were correlated to PII in terms of protein abundance, by plotting their correlation against PII on the x-axis and the slope of this correlation on the y-axis (Figure 6B, numerical data in Supplementary Table S14). These analyses show that *Rhizobium* sp. *Root491* shows the highest nitrogen stress response under these nitrogen treatments, with all three amino acid treatments leading to dramatic increases in the abundance of the PII protein, and also with many more proteins positively correlated to PII abundance in Rhizobium sp. Root491 compared to the other two strains. Looking at the identity of proteins whose abundance was correlated to PII in *Rhizobium* sp. *Root491*, we see that 10 proteins controlled by the exo operon that conduct the synthesis and export of extracellular polysaccharides are positively correlated to PII abundance (Supplementary Table S14). Analogous findings have been reported via genetic manipulation of *V. vulnificus* and *S. meliloti*, with knockout of nitrogen stress response elements NtrC and NtrX resulting in reduced production of extracellular polysaccharides (35, 36). In *Pseudomonas* sp. *Root9*, the data point that exhibits a strong negative correlation to PII is an NADP-dependent glutamate dehydrogenase (Supplementary Table S14), previously shown to be a target of NtrC-driven transcriptional repression in *P. putida* (37).

**Figure 6:**
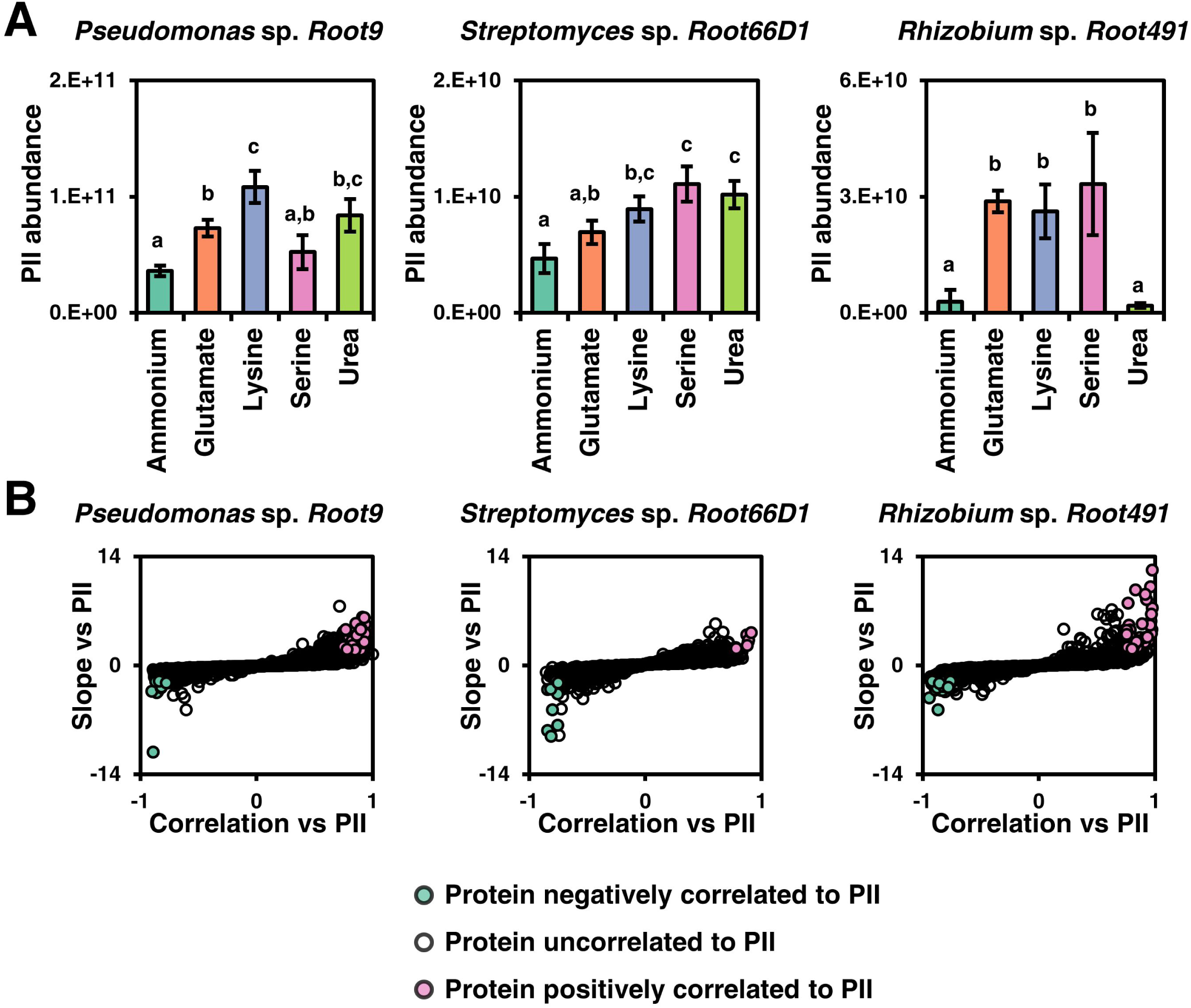
Investigating proteins correlated to the abundance of nitrogen stress response component PII. A: Abundance of the PII protein across five nitrogen treatments in three rhizosphere bacterial strains. Different letters above data series indicate p<0.05 following two-way ANOVA and Tukey’s HSD test. B: Plots to highlight proteins that are positively or negatively correlated to PII according to their abundance values across five nitrogen treatments. Y-displays the slope of the linear fit (z-score normalised) between protein abundance versus the abundance of PII protein, and X-axis displays correlation between protein abundance versus PII abundance. If a protein has a correlation higher than 0.75 and a slope higher than 2, it is deemed positively correlated, whereas if a protein has a correlation lower than 0.75 and a slope lower than −2, it is deemed negatively correlated to PII.

## Discussion

Differential nitrogen treatments are a classical experimental manipulation in microbiology, but the majority of molecular knowledge about bacterial nitrogen metabolism has been acquired in *E. coli* (14). To deepen our knowledge of nitrogen metabolism in the rhizosphere microbiome, this study analyses nitrogen substrate utilisation in three taxonomically diverse bacterial strains previously isolated from field-grown Arabidopsis roots (20). The three strains represent taxa that are consistently detected as core members of the plant microbiome: Pseudomonas, Streptomyces and Rhizobium (21). Using label-free quantitative proteomics, we document hundreds of proteins in each strain that exhibit differential abundance values between nitrogen sources. To enable protein-level comparisons between these taxonomically diverse strains, we integrate the identified proteins using KEGG Orthologues, and map the differential expression of orthologous proteins onto metabolic maps to determine which specific metabolic pathways are modulated by nitrogen source at the protein level. We also determine novel proteins linked to the nitrogen stress response in these three strains, by investigating which proteins display abundance values that are positively and negatively correlated to the PII signal transduction protein. Furthermore, we integrate experimental data with computational models, using the EnsembleFBA method to test how accurately metabolic phenotypes can be computationally predicted from a minimal set of experimental data. Our results show that the three strains exhibit diverse metabolic responses to different nitrogen nutrition regimes, with a summary of key results presented in Supplementary Table S15.

One noticeable observation in our quantitative proteomic dataset is that the three different bacterial strains exhibit widely divergent protein-level responses to the same nitrogen source. This is best illustrated in the pairwise comparisons of protein composition between two nitrogen sources, which yield dramatic variation in the number of differentially expressed proteins across the three strains. For example, the ammonium versus serine pairwise comparison resulted in only eight DEPs for *Pseudomonas* sp. *Root9*, but 74 DEPs in *Streptomyces* sp. *Root66D1* and 100 DEPs in *Rhizobium* sp. *Root491*. Reciprocal responses were observed for pairwise comparisons between ammonium versus lysine nutrition, which elicited widespread alterations to the proteome of *Pseudomonas* sp. *Root9* but relatively fewer protein-level changes in the other two studied strains. One potential explanation for this difference is that the minimally responsive strains induce enzymes that can convert these different nitrogen sources into ammonium via relatively simple metabolic pathways. For *Pseudomonas* sp. *Root9*, one of the few proteins induced under serine nutrition is serine dehydratase, which yields ammonium in one enzymatic step, along with pyruvate that can be quickly assimilated in the TCA cycle. In contrast, the other two studied strains exhibited no upregulation of their serine dehydratase proteins under serine nutrition, potentially indicating the assimilated serine must be distributed through multiple elements of the metabolic network requiring a wider modulation of protein expression. For lysine, our proteomics data indicate that lysine degradation in *Pseudomonas* sp. *Root9* proceeds via the δ-aminovalerate pathway, whereas *Rhizobium* sp. *Root491* appears to utilise the saccharopine pathway of lysine degradation. Although both of these pathways yield relatively similar products and contain a similar number of enzymatic steps, our data indicate that the operation of the δ-aminovalerate pathway in *Pseudomonas* sp. *Root9* could require a dramatic remodelling of cellular protein composition, and a much longer lag phase before cell proliferation can begin. In contrast, *Rhizobium* sp. *Root491* shows almost identical growth curves on lysine and ammonium, and the relatively small set of proteins modulated by lysine nutrition contains a high proportion of transporters. Although both strains exhibit generalistic nitrogen substrate preferences, the contrasting proteomic impact of lysine nutrition indicates that *Pseudomonas* sp. *Root9* metabolises diverse substrates by adapting its highly flexible metabolic network, whereas *Rhizobium* sp. *Root491* utilises different transport mechanisms to assimilate diverse nitrogen sources into a relatively stable metabolic network.

There is a longstanding appreciation that amino acids play a significant role in the nutrition of rhizosphere bacterial strains (38). Amino acids are an important component of the soil nitrogen cycle, derived from diverse sources such as depolymerisation of soil bound protein and also from plant rhizodeposition (18). Microbial metabolism of amino acids in the rhizosphere is related to plant productivity, because microbial mineralisation of organic nitrogen can boost plant nutrition (39), while the microbial uptake of amino acids is one mechanism used by plants to recruit specific strains into the rhizosphere microbiome (40). The data presented here could potentially assist future efforts to manipulate the rhizosphere microbiome for altered metabolism of amino acids. For instance, our data in *Pseudomonas* sp. *Root9* implicate serine dehydratase as an important protein for degradation of serine, and measurements in *Rhizobium* sp. *Root491* position saccharopine dehydrogenase as important for degradation of lysine. Perhaps bacterial strains with high activities of these two enzymes could be recruited to the rhizosphere to promote faster rates of amino acid mineralisation. In *Rhizobium* sp. *Root491*, we document that this strain grows quickly on three chemically diverse amino acids, and also that dozens of ABC transporter proteins exhibit altered abundance values under amino acid nutrition. Previous work in *E. coli* has shown amino acids such as glutamate and arginine serve as poor sole nitrogen sources for enteric bacteria, with this phenotype being underpinned by slow rates of amino acid transport and catabolism (41). Perhaps the protein network that undertakes amino acid transport and catabolism in *Rhizobium* sp. *Root491* could serve as a template for engineering other bacterial strains to grow rapidly on amino acids as a sole nitrogen source. In *Streptomyces* sp. *Root66D1*, amino acid nutrition results in upregulation of dozens of proteins, but very few of these are classically recognised as being involved in amino acid degradation. Compared to other bacterial taxa, there is generally less knowledge about nitrogen metabolism in Gram-positive Streptomyces (16), so the uncharacterised proteins shown to be differentially expressed under amino acid nutrition in *Streptomyces* sp. *Root66D1* could be targets for future studies investigating their biochemical function.

Urea is the most widely applied agricultural fertiliser globally, but plant nutrition experiments show that urea is a relatively poor sole nitrogen source for plant growth (42). Although plants can uptake urea to some degree, a large proportion of the nitrogen delivered via urea fertilisers must first undergo hydrolysis by microbial metabolism before it can subsequently contribute to plant nutrition (43). Therefore, urea metabolism in the rhizosphere microbiome is a potential target for improving agricultural nitrogen use efficiency. In our work, we show that all three tested strains can grow rapidly on urea as a sole nitrogen source. However, the proteomic impact of urea nutrition differed widely between the three strains, with *Streptomyces* sp. *Root66D1* and *Rhizobium* sp. *Root491* both showing zero proteins that were differentially expressed between ammonium versus urea treatment, whereas this comparison in *Pseudomonas* sp. *Root9* elicited 126 differentially expressed proteins. The urease enzyme that converts urea to ammonium is required under normal conditions for catabolism of purine and arginine, and is increasingly expressed under nitrogen stress as a nutrient salvage mechanism. In our dataset, all three strains exhibit high expression of urease subunits under all conditions tested, and our investigations of the nitrogen stress response showed that many urease subunits are tightly correlated to PII expression. For all three strains, we see that at least one amino acid treatment actually elicits a higher urease expression compared to urea nutrition. This suggests that urease abundance is not the limiting factor for utilisation of urea as a sole nitrogen source, and that other mechanisms may explain urea-induced proteome remodelling in *Pseudomonas* sp. *Root9*. Inspecting the data, we see many transporter proteins are differentially expressed in *Pseudomonas* sp. *Root9* under urea versus ammonium nutrition, which may be involved in urea uptake or the excretion of urea-derived waste products. In comparison, the transport machineries of both *Streptomyces* sp. *Root66D1* and *Rhizobium* sp. *Root491* seem to already be primed for urea uptake when cultivated on ammonium. Future studies could investigate how to optimally coordinate urea transport and metabolism between plants and rhizosphere microbes to deliver higher nitrogen use efficiency from urea fertilisers.

Many microbial strains have been labelled as plant growth promoting, but there is relatively little knowledge about the genes and mechanisms that underpin this trait (44). In previous work, *Rhizobium* sp. *Root491* was characterised as a plant growth promoting bacterium by its ability to increase Arabidopsis root length in co-cultivation experiments (45). Furthermore, exometabolomics profiling has shown that *Rhizobium* sp. *Root491* can consume a wide variety of plant-derived metabolites as carbon substrates (46). Here, we show that *Rhizobium* sp. *Root491* exhibits fast growth on a variety of nitrogen sources, that its set of ABC transporters exhibit differential abundance values in response to nitrogen source, and also that amino acid nutrition induces the expression of multiple proteins involved in the production of extracellular polysaccharides. When combined with previous observations of *Rhizobium* sp. *Root491*, we can begin to characterise the functional traits possessed by this strain that contribute to plant growth promotion, such as: recruitment to the rhizosphere via the consumption of plant root metabolites, adherence to the root surface via biofilm production in the presence of plant-derived amino acids, and the potential for mineralisation of diverse nitrogen molecules to fuel plant nutrition. Potentially, future studies could predict whether other rhizosphere strains can also promote plant growth via similar mechanisms, by investigating genetic similarities with *Rhizobium* sp. *Root491*. Also, future work could investigate whether plant genotypes differ in their ability to attract growth-promoting strains to the rhizosphere, and how to design synthetic microbial communities that combine multiple growth-promoting strains.

There is increasing interest in combining experimental and computational approaches to analyse microbial metabolism, with the long-term goal of quantitatively predicting the behaviour of microbial communities (47). Metabolic modelling is rapidly progressing as a powerful computational tool to explore the metabolic capacities of bacteria. However, the main limitation that prevents modelling approaches from being applied to diverse bacterial strains is the need to obtain a highly curated genome-scale metabolic model for each strain of interest. This process of model curation still requires a significant amount of manual inspection and relies heavily on accurate genome annotation (25). In the present study, we used EnsembleFBA (26) to produce metabolic models for three diverse bacterial strains using a minimal set of experimental information. We compared the derived models versus experimental data by assessing how accurately they can predict growth phenotypes and proteome remodelling across different nitrogen sources. This showed that EnsembleFBA gives relatively accurate predictions of nitrogen substrate utilisation, with binary phenotypes (growth versus no growth) correctly predicted in around 80% of cases. However, there was only an intermediate correlation between the proxy values of metabolic activity predicted by the model versus the experimentally acquired measurements (r^2^: 0.23-0.50), and a relatively poor concordance between the differential fluxes predicted by the model versus the differentially expressed proteins identified via proteomics. We present two potential interpretations for these inaccurate predictions. First, there is no straightforward relationship between enzymatic flux and protein abundance, because the catalysis rate of many enzymes is not only regulated via abundance but also by other factors including post-translational modifications, allosteric regulators or the relative concentrations of substrates and products (48). Second, our models used the same biomass definition that Biggs and Papin used for their EnsembleFBA analyses of *Pseudomonas* and *Streptococcus* (26). Although efforts have been made to define a general biomass composition for bacteria (49), inaccuracies of this definition can decrease the predictive power of metabolic models. Therefore, one potential pathway to improve model accuracy would involve measuring the biomass composition for all genotypes and treatments under study. Despite these limitations, our work shows that EnsembleFBA shows strong potential for predicting nitrogen substrate utilisation across diverse bacterial strains, using minimal experimental data and requiring no manual curation of the model.

Manipulating the rhizosphere microbiome is one proposed solution to reduce the application of synthetic chemicals in agriculture, particularly mineral nitrogen fertilisers (3). Plant microbiome research is being advanced by the collection of thousands of genomically sequenced bacterial strains isolated from the plant host (20, 21). A current research priority is to characterise the functional traits encoded by plant-associated microbial strains, in order to rationally design synthetic microbial communities that can promote plant growth and health (50). Here we analyse nitrogen metabolism in three bacterial strains previously isolated from field-grown Arabidopsis roots using a combination of experimental and computational approaches. From the growth analyses, it is evident that all three strains can utilise a large and similar set of nitrogen substrates. However, proteomic measurements showed that the strains deploy different metabolic strategies to utilise specific nitrogen sources. One diverging trait appears to be their degree of proteomic flexibility, with *Pseudomonas* sp. *Root9* utilising lysine via widespread protein-level alterations to its flexible metabolic network. In contrast, *Rhizobium* sp. *Root491* shows relatively stable proteome composition across diverse nitrogen sources, characterised by minimal alterations to central metabolism but differential abundance of many transport proteins. In addition, we document a large set of functionally uncharacterised proteins that display differential abundance values in response to nitrogen source, with functional annotations being particularly unclear in Gram-positive *Streptomyces* sp. *Root66D1*. These proteins are potentially important for nitrogen metabolism in the rhizosphere, and could be the targets of future functional study. Our results could inform the selection of high-performing strains in synthetic microbial communities designed to mediate plant nitrogen nutrition under lower inputs of mineral fertilisers.

## Materials and methods

### Bacterial strains

Bacterial strains used in this study were *Pseudomonas* sp. *Root9* (NCBI Taxonomy ID: 1736604), *Streptomyces* sp. *Root66D1* (NCBI Taxonomy ID: 1736582) and *Rhizobium* sp. *Root491* (NCBI Taxonomy ID: 1736548), all isolated from field-grown Arabidopsis roots (20), and provided by Paul Schulze-Lefert, MPIPZ Cologne.

### Bacterial pre-cultivation and harvest

Bacterial strains were pre-cultivated by streaking glycerol stocks onto TSA plates (0.5× TSB, 1.2% Agar), and incubating at 28° C for 24 hours. Single colonies were picked from plates and inoculated into TSB medium (0.5× TSB), and incubated for 24 h at 28° C with 200 rpm shaking. Next, cells were harvested by centrifuging 800 μL of culture at 5,000× g for 2 min at RT. These cells were then rinsed 3× in sterile 10 mM MgCl_2_, and resuspended at a final OD_600_ of 1.0 in sterile 10 mM MgCl_2_.

### Phenotype microarrays

For phenotype microarrays using PM3B (Biolog), 12 ml of inoculant was prepared comprising 10 mL of 1.2× IF-0 (Biolog), 1.2 mL of 500 mM glucose, 600 uL of bacterial suspension (as prepared above), 120 uL of Redox Dye D (Biolog) and 80 uL of sterile water. Next, 100 uL of this inoculant (starting OD_600_ of 0.05) was loaded into each well of the phenotype microarray, which was transferred to a plate reader (Tecan Infinite Pro 100) and incubated at 28° C for 72 h with shaking (30 sec continuous orbital shaking followed by 9:30 min stationary, shaking amplitude 3 mm). Tetrazolium reduction at A_590_ was measured once per 10 min cycle, without correcting for path length, and derived curves were fitted to a logistic equation using the Growthcurver program (51). For each well in every assay, background was subtracted by subtracting the value of the negative control (well A1) from each time point. In our hands, guanosine (well F7) gave a very high background reading and was excluded from the analysis. Wells were considered growth-positive if the carrying capacity (k) of the logistic fit was greater than A_590_ of 0.1 in at least two of the three independent biological replicates. Next, area under the curve (AUC) values for all growth-positive wells were z-score normalised within each strain, and the average value of the three replicate assays was calculated. These averaged z-score values were divided into quartiles, so data presented in Fig 1 represent five possible growth intensities, ranging from 0 (no growth) to 4 (highest AUC quartile).

### Metabolic models and computational simulations

The EnsembleFBA workflow from Biggs and Papin (26) was adapted to analyse the three studied bacterial strains. Scripts were implemented either in Matlab (Mathworks) as the original code, or adapted for Python (Python Software Foundation). Briefly, genomes were downloaded from NCBI (52) and uploaded to KBase (25), where genome re-annotation and draft metabolic model reconstruction was performed. Outputted draft networks were downloaded and used as inputs for the EnsembleFBA workflow. Also inputted to Ensemble FBA were the composition of the Biolog media, and the experimentally derived growth matrices obtained from PM3B phenotype microarray. Next, 50 metabolic networks were generated for each strain, with each network being trained on 26 nitrogen substrates that supported growth and 11 nitrogen substrates that didn’t support growth, in order to perform positive and negative gapfilling. Compounds present on the phenotype microarray but not found in the ModelSEED database (24) were excluded, and a second set of simulations excluding the five N-sources used for proteomics experiments were also obtained for unbiased integration with the proteomics datasets. To evaluate the performance of EnsembleFBA for predicting growth on the different N-sources, its accuracy, precision and recall were compared to randomly generated predictions, after masking the conditions used to gapfill the individual networks to avoid bias. Metabolic activity on a given nitrogen source was estimated as the average growth rate obtained with EnsembleFBA, and weighted according to the fraction of networks in the ensemble that predicted growth. Metabolic fluxes through specific reactions were estimated by averaging the reaction flux for each reaction across all the networks in the ensemble, and weighted according to the fraction of networks where the reaction was active. To visualise up- or down-regulated metabolic fluxes in metabolic pathway maps, metabolic fluxes obtained by simulating growth on Glutamate, Serine or Lysine were compared versus Ammonium, and filtered for reactions with log2 fold change greater than 1.

### Cultivation on individual N-sources for growth assays and proteomic analysis

For growth assays on individual N-sources, media were based on M9 formulation (53), with nutrient concentrations of: 50 mM glucose, 24 mM Na_2_HPO_4_, 11 mM KH_2_PO_4_, 4 mM NaCl, 350 μM MgSO_4_, 100 μM CaCl_2_, 50 μM Fe-EDTA, 50 μM H_3_BO_3_, 10 μM MnCl_2_, 1.75 μM ZnCl_2_, 1 uM KI, 800 nM Na_2_MoO_4_, 500 nM CuCl_2_, 100 nM CoCl_2_. To this, one nitrogen source was added at 5 mM elemental-N (ie: 5 mM of ammonium, glutamate and serine, or 2.5 mM of urea and lysine). For growth assays, 20 μL of bacterial suspension (as prepared above) was inoculated into 380 μL of growth medium (starting OD_600_ of 0.05), in individual wells of a sterile 48-well plate (Corning). These plates were then transferred to a plate reader (Tecan Infinite Pro 100) and incubated at 28° C for 48 h with shaking (3 min continuous orbital shaking followed by 7 min stationary, shaking amplitude 3 mm). Culture density at OD_600_ was measured once per 10 min cycle, without correcting for path length. To obtain quantitative growth metrics, a logistic equation was fitted to measured growth curves using the Growthcurver program (51). To collect samples for proteomics, cultivation was identical, except that bacterial cells were harvested during the exponential growth phase. Harvest involved pooling of four duplicate wells (total of 1.6 mL culture), followed by centrifugation at 10,000× g for 3 min at 4° C. Supernatant was discarded, and cell pellets were rinsed twice with 900 uL of 4° C PBS via centrifugation at 10,000x g for 3 min at 4° C. Rinsed cell pellets were then flash-frozen and stored at −80° C.

### Proteomic sample preparation

Cellular protein was extracted using protocols modified from Tanca et al (54) as well as Wessel and Flugge (55). To frozen cell pellets, 250 uL of lysis buffer (5% SDS, 100 mM DTT, 100 mM Tris pH 7.5) was added, along with ∼100 uL of acid-washed glass beads (1 mm diameter). Samples were then incubated for 10 min on an orbital mixer at 95° C with 1500 rpm shaking, then at −80° C for 10 min, then bead-beaten (Bead Ruptor 24, Omni International) at 5 ms-1 for 10 min. Next, samples were again incubated at −80° C for 10 min, then again incubated for 10 min on an orbital mixer at 95° C with 1500 rpm shaking, then again bead-beaten at 5 ms-1 for 10 min. Finally, samples were centrifuged at 20,000x g for 10 min at RT, and 200 uL of supernatant was transferred to a new tube. Protein was then precipitated via the addition of 800 uL MeOH, 500 uL H_2_O, and 200 uL chloroform followed by centrifugation at 10,000x g for 5 min at 4° C. The upper aqueous phase was removed and discarded, then 700 uL MeOH was added to the lower organic phase and samples were centrifuged at 20,0000x g for 10 min at 4° C. Protein pellets were then rinsed twice with −20° C acetone via centriguation at 20,0000x g for 10 min at 4d C, before being air-dried at RT for 15 min. Dried protein pellets were then stored at −80° C. To solubilise protein pellets, 40 uL of solubilisation buffer (8 M urea, 50 mM TEAB, 5 mM DTT) was added, and samples were incubated on an orbital mixer at 28° C for 1 h with 350 rpm mixing. Next, CAA was added to a final concentration of 30 mM, and samples were incubated on an orbital mixer at 28° C for 30 min with 350 rpm mixing in darkness. To quantify protein concentration, an aliquot of the protein extract was taken and diluted 1:8 in water, then a Bradford assay was performed on the diluted protein samples using BSA as standard. Next, 40 ug of protein extract was transferred to a new tube and incubated with 0.8 ug Lys-C for 2 h at 37d C with 350 rpm shaking. Samples were then diluted 1:8 in TEAB, 0.8 ug of trypsin was added, and samples were incubated overnight at 37° C. Next day, samples were acidified by adding formic acid to a final concentration of 1%. Peptides were then cleaned up via SPE using SDB-RP stage tips. Following elution from stage tips, peptides were dried down in a vacuum centrifuge and stored at −80° C.

### Mass spectrometry

Digested peptides were analysed on a QExactive Plus mass spectrometer (Thermo Scientific) coupled to an EASY nLC 1000 UPLC (Thermo Scientific). Dried peptides were resolubilised in solvent A (0.1% formic acid), and loaded onto an in-house packed C18 column (50 cm × 75 µm I.D., filled with 2.7 µm Poroshell 120, (Agilent)). Following loading, samples were eluted from the C18 column with solvent B (0.1% formic acid in 80% acetonitrile) using a 2.5 h gradient, comprising: linear increase from 4-27% B over 120 min, 27-50% B over 19 min, followed by column washing and equilibration. Flow rate was at 250 nL/min. Data-dependent acquisition was used to acquire MS/MS data, whereby the 10 most abundant ions (charges 2-5) in the survey spectrum were subjected to HCD fragmentation. MS scans were acquired from 300 to 1750 m/z at a resolution of 70,000, while MS/MS scans were acquired at a resolution of 17,500. Following fragmentation, precursor ions were dynamically excluded for 25 s.

### Label-free protein quantification

Label-free quantification of protein abundance was conducted with MaxQuant v1.5.3.8 (56). Acquired MS/MS spectra were searched against FASTA protein sequences for the three studied bacterial strains, obtained from IMG (57). Sequences of common contaminant proteins were also included in the search database. Protein FDR and PSM FDR were set to 0.01%. Minimum peptide length was seven amino acids, cysteine carbamidomethylation was set as a fixed modification, while methionine oxidation and protein N-terminal acetylation were set as variable modifications.

### Statistical analysis of proteomic data

To determine proteins that exhibited significantly different abundance between N-treatments, a statistical threshold was imposed where the MaxQuant LFQ values must differ by log_2_FC > 1 and BH-p-value <0.05. To determine the abundance of Kegg Orthologues (KOs) across bacterial strains and N-treatments, KOs annotated to proteins via IMG were matched across bacterial strains. Data were filtered to contain only the 495 KOs that were observed in at least three replicates across all five treatments in all three strains. In instances where a single strain had multiple proteins matching the same KO, the protein with the highest average MaxQuant LFQ value across all samples was taken as the representative KO for that strain. To determine the KEGG pathways that were significantly modulated at the protein abundance level between nitrogen treatments, KOs annotated onto proteins via IMG were mapped against KEGG pathways using KEGG-REST, and Fisher’s exact test was used to generate a single p-value for each KEGG pathway by combining the individual BH-p-values for all constituent proteins mapped to that pathway. Pathways were only analysed when at least three representative proteins were detected for a single strain across all five nitrogen treatments, and pathways associated with non-bacterial processes were discarded.

## Data availability

All LC-MS/MS files and MaxQuant outputs have been uploaded to ProteomXchange and can be accessed via PRIDE (URL: https://www.ebi.ac.uk/pride/archive/, Accession: PXD011436, Username, Password: IG63IYVi). Details of the EnsembleFBA workflow are available at: https://github.com/asuccurro/ensembleFBA, and the KBase narrative is available at https://narrative.kbase.us/narrative/ws.37070.obj.1. Interactive maps of metabolic pathways modulated between amino acid treatments can be viewed at: https://pathways.embl.de/shared/rjacoby.

## Acknowledgements

We thank Paul Schulze-Lefert (Max Planck Institute for Plant Breeding, Cologne) for providing bacterial strains, as well as Christian Frese and Corinna Klein (Proteomics Core Facility Cologne, University of Cologne) for conducting proteomics measurements. RPJ is funded by a Humboldt Research Fellowship, and previously by the Horizon 2020 Marie Curie Sklodowska Action project 705808 – PINBAC. Research in SK’s lab is funded by the Deutsche Forschungsgemeinschaft (DFG) under Germany’s Excellence Strategy – EXC 2048/1 – project 390686111.

## Supplementary Figure captions

Supplementary Figure S1: Nitrogen substrate preferences of three rhizosphere bacterial strains assessed via Phenotype Microarray and EnsembleFBA. Displayed here are metabolic activity values for 94 nitrogen substrates measured via phenotype microarray and 81 nitrogen substrates predicted via EnsembleFBA. White boxes indicate no metabolic activity, while boxes with darker shades correspond to higher metabolic activity, either measured via Phenotype Microarray (pink) or predicted via EnsembleFBA (green). Metabolic activity values were z-score normalised within each strain.

Supplementary Figure S2: Comparison of measured versus predicted nitrogen substrate utilisation for three rhizosphere bacterial strains. A: Venn diagram showing nitrogen substrate utilisation for three bacterial strains as measured using phenotype microarray. B, C and E: Plots showing the correlation between predicted metabolic activity (EnsembleFBA) versus measured metabolic activity (EnsembleFBA) for 81 nitrogen substrates across three bacterial strains. D: Venn diagram nitrogen substrate utilisation for three rhizosphere bacterial strains as predicted using EnsembleFBA.

Supplementary Figure S3: Determination of ammonium concentration where nitrogen is the yield limiting nutrient in batch culture. Three rhizosphere bacterial strains were cultivated on media containing different concentrations of NH_4_Cl, and OD_600_ was logged every 10 min in a plate reader. Logistic growth equations were fitted to derived growth curves, and in these graphs the carrying capacity (k) of the logistic fits is plotted against NH_4_Cl concentration. There is a linear relationship between k and NH_4_Cl concentration until around 10 mM, indicating that N is the yield-limiting nutrient at these concentrations. Therefore, nitrogen was supplied at 5 mM N for all experiments.

Supplementary Figure S4: Volcano plots of proteomic data for *Pseudomonas* sp. *Root9* grown on five different nitrogen sources. For proteins that were detected in three or more replicates in both treatments, the Y-axis shows –log10 of the Benjamini-Hochberg p-value, while X-axis shows the log2 fold change. Proteins with a log2 fold change ≥ 1 and Benjamini-Hochberg p-value ≤ 0.05 are deemed differentially expressed and rendered in colour. In total, 10 comparisons were performed, A: Ammonium vs Glutamate, B: Ammonium vs Lysine, C: Ammonium vs Serine, D: Ammonium vs Urea, E: Glutamate vs Lysine, F: Glutamate vs Serine, G: Glutamate vs Urea, H: Lysine vs Serine, I: Lysine vs Urea, J: Serine vs Urea:

Supplementary Figure S5: Volcano plots of proteomic data for *Streptomyces* sp. *Root66D1* grown on five different nitrogen sources. For proteins that were detected in three or more replicates in both treatments, the Y-axis shows –log10 of the Benjamini-Hochberg p-value, while X-axis shows the log2 fold change. Proteins with a log2 fold change ≥ 1 and Benjamini-Hochberg p-value ≤ 0.05 are deemed differentially expressed and rendered in colour. In total, 10 comparisons were performed, A: Ammonium vs Glutamate, B: Ammonium vs Lysine, C: Ammonium vs Serine, D: Ammonium vs Urea, E: Glutamate vs Lysine, F: Glutamate vs Serine, G: Glutamate vs Urea, H: Lysine vs Serine, I: Lysine vs Urea, J: Serine vs Urea:

Supplementary Figure S6: Volcano plots of proteomic data for *Rhizobium* sp. *Root491* grown on five different nitrogen sources. For proteins that were detected in three or more replicates in both treatments, the Y-axis shows –log10 of the Benjamini-Hochberg p-value, while X-axis shows the log2 fold change. Proteins with a log2 fold change ≥ 1 and Benjamini-Hochberg p-value ≤ 0.05 are deemed differentially expressed and rendered in colour. In total, 10 comparisons were performed, A: Ammonium vs Glutamate, B: Ammonium vs Lysine, C: Ammonium vs Serine, D: Ammonium vs Urea, E: Glutamate vs Lysine, F: Glutamate vs Serine, G: Glutamate vs Urea, H: Lysine vs Serine, I: Lysine vs Urea, J: Serine vs Urea:

Supplementary Figure S7: Differentially expressed proteins mapped onto metabolic pathways for three rhizosphere bacterial strains cultivated on amino acids as the sole nitrogen source. Protein abundance data from each of the three amino acid treatments (glutamate, lysine and serine) is compared against the ammonium control. For proteins that were detected in three or more replicates in both treatments, Kegg orthologs annotated to proteins via IMG were matched to the ‘Metabolic Pathways’ map provided via the Interactive Pathways Explorer v3. KOs matching proteins with a log2 fold change ≥ 1 and Benjamini-Hochberg p-value ≤ 0.05 are deemed differentially expressed and rendered in colour. KOs matching proteins that were not differentially expressed are rendered in grey. Interactive maps can be viewed at: https://pathways.embl.de/shared/rjacoby.

Supplementary Figure S8: Differentially expressed fluxes mapped onto metabolic pathways for three rhizosphere bacterial strains cultivated on amino acids as the sole nitrogen source. Modelled flux from each of the three amino acid treatments (glutamate, lysine and serine) is compared against the ammonium control. Kegg reactions annotated via KBase were matched to the ‘Metabolic Pathways’ map provided via the Interactive Pathways Explorer v3. Reaction fluxes with a log2 fold change ≥ 1 are rendered in colour. Fluxes that were not different expressed are rendered in grey. Interactive maps can be viewed at: https://pathways.embl.de/shared/asuccurro.

## Supplementary Table captions

Supplementary Table S1: Nitrogen substrate utilisation of three rhizosphere bacterial strains assessed by phenotype microarray measurement for 94 nitrogen sources and EnsembleFBA prediction for 81 nitrogen sources.

Supplementary Table S2: Assessment of concordance between EnsembleFBA predictions and experimental measurements of nitrogen substrate utilisation across three rhizosphere bacterial strains.

Supplementary Table S3: Growth curve metrics for three bacterial strains cultivated on five nitrogen substrates.

Supplementary Table S4: Protein abundance information acquired from label-free proteomic profiling of three rhizosphere bacterial strains cultivated on five nitrogen sources.

Supplementary Table S5: Protein abundance values mapped to KEGG Orthologues for three rhizosphere bacterial strains cultivated on five nitrogen sources.

Supplementary Table S6: Determination of KEGG pathways with differentially expressed proteins calculated via Fisher’s exact test of protein abundance values.

Supplementary Tables S7-S8: Metabolic model parameters for *Pseudomonas* sp. *Root9* generated by KBase.

Supplementary Tables S9-S10: Metabolic model parameters for *Streptomyces* sp. *Root66D1* generated by KBase.

Supplementary Tables S11-S12: Metabolic model parameters for *Rhizobium* sp. *Root491* generated by KBase.

Supplementary Table S13: Biomass components used for models generated by EnsembleFBA, taken from Biggs and Papin (2017).

Supplementary Table S14: Determination of proteins that are correlated to the PII component of the nitrogen stress response in three rhizosphere bacterial strains cultivated on five nitrogen sources.

Supplementary Table S15: Summary of key results obtained regarding nitrogen metabolism in the three rhizosphere bacterial strains studied.

